# Visualization of purine and pyrimidine *de novo* synthesis and salvage pathway activity in single-cell using Carbon Isotope Imaging and Spectral Tracing (CIIST)

**DOI:** 10.1101/2025.02.28.640723

**Authors:** Jiro Karlo, Tejas Ajit Mhaiskar, Hrishikesh Ravindra Karande, S P Singh

## Abstract

Nitrogenous bases, namely purine and pyrimidine, and their derivatives are key metabolites for the growth and division of cells as they are involved in the storage of genetic information, protein synthesis, energy carrier molecules, and many other metabolic processes. Here we report a single-cell Raman imaging technique for nitrogenous base pathway mapping in prokaryotic and eukaryotic microbial systems via carbon isotope imaging and spectral tracing (CIIST). This method helps in visualizing the turnover dynamics of *de novo* synthesized purine and pyrimidine nitrogenous bases at the sub-cellular level over time. The enrichment of carbon isotope (carbon-12 or carbon-13) in the nitrogenous base generates Raman peaks at different positions. CIIST can also help in visualizing the salvage pathway activity by identifying new exogenously transported purine and pyrimidine into the cell. CIIST can also be used for generating spatial maps for quasi-quantitative imaging of nitrogenous base turnover in cells. Overall findings provides support for the prospective utility of CIIST technique as a highly effective tool for spatiotemporal and multiplex biomolecular analysis of nitrogenous base metabolism with unmatched spatial resolution in a single cell.

## Introduction

The growth and division of microbes are highly dependent on derivatives of nitrogenous bases such as adenosine triphosphate (ATP), guanosine triphosphate (GTP) and nucleic acids such as deoxyribose nucleic acid (DNA) and ribose nucleic acid (RNA). Purines (adenine and guanine) and pyrimidines (cytosine, thymine and uracil) are small nitrogen-containing organic bases (nitrogenous bases) with double and single heterocyclic rings. It is mostly found attached to a sugar group known as nucleoside or attached to sugar phosphates known as nucleotides.[1] *De novo* pathway is the synthesis of these molecules from scratch and the salvage pathway is the reutilization from existing or external sources. ATP and GTP are known as the major energy carrier molecules, cyclic di-GMP acts as a signalling molecule, DNA stores the genetic information and RNA plays a major role in protein synthesis. Analysing nitrogenous bases metabolic pathways is critical for understanding microbial colonization, survival, and virulence. For instance, the synthesis of purine and pyrimidines is required for *Escherichia coli* to colonize and persist in the mouse intestine.[2] Additionally, the connection between the nucleotide biosynthesis pathway and the pathogenesis of bacteria such as *Staphylococcus aureus* has been reported previously.[3] Therefore, targeting and monitoring nucleotide biosynthesis can act as an antivirulence strategy in existing antimicrobial therapies. In higher organisms, many metabolic disorders such as orotic aciduria, adenosine deaminase deficiency, purine nucleoside phosphorylase deficiency, hyperuricemic metabolic disorders are linked to dysfucntioning of purine and pyrimidine metabolism.[4]

Purine and pyrimidine can either be synthesized, recycled and/or acquired exogenously which is categorised as: (i) *de novo* pathway and (ii) salvage pathway.[5–10] These nitrogenous bases can be analysed through existing gold-standard mass spectrometry-based methods with high sensitivity.[11] However, the traditional methods are inherently destructive. Towards this unmet goal of non-destructive and minimal invasive visualization of the nascent nitrogenous base at the single cell level, herein we combined the Raman imaging with carbon isotope tracing microbial methodology. Raman spectroscopy coupled with different and stable isotopes of hydrogen, carbon, and nitrogen for detecting metabolites has been studied previously.[12–20] The isotopic substitution in the metabolites leads to a shift in spectral peaks due to the relation between the reduced mass and the frequency; if the heavier isotope substitutes the lighter isotope the shift occurs towards a lower wavenumber known as “red-shift” and if the lighter isotope substitutes the heavier isotope, the shift occurs towards higher wavenumber known as “blue-shift”.[15, 17, 18, 21] Taking merit of this exquisite sensitivity of Raman spectroscopy for isotope substitution, here we introduce a microbial nitrogenous pathway activity imaging methodology known as Carbon Isotope Imaging and Spectral Tracing (CIIST). CIIST involves two carbon isotopes and tracks the enrichment of the carbon isotopes acting as nascent nitrogenous base tracers. Raman-based CIIST method tracks and maps the nitrogenous bases in their physiological environment in an intact single-cell level with minimal reagent requirement and in a non-destructive way.

This study’s aims are three-fold: First, to map the pool of newly *de novo* synthesized nitrogenous bases in bacteria and yeast both qualitatively and quasi-quantitatively. Second, to spectral trace the cell-to-cell heterogeneity of *de novo* synthesis pathway activity at single cell level using point spectroscopy, and third to map the salvage pathway activity in bacteria and yeast using CIIST. For this study, we administered ^12^C glucose as the only carbon source to the ^13^C enriched microbes and traced the shift in the peak corresponding to nitrogenous bases in the Raman spectra of cells. Targeting the blue shift, we have mapped the nascent nitrogenous bases with high spatial information in a single microbial cell.

## Methods

### Microbial culture condition and carbon isotope cell preparation

Microbial cells used in this study were *Escherichia coli* (bacteria) and *Saccharomyces cerevisiae* (yeast). For ^13^C isotope bacterial cells preparation, ^12^C *E. coli* cells were grown in carbon source-free M9 minimal medium (Sigma) supplemented with 5 g/L uniform ^13^C glucose (Cambridge isotope laboratories) and incubated overnight in a rotational shaker at 180 rpm and 37°C incubation temperature. For the preparation of ^13^C isotope yeast cells, ^12^C *S. cerevisiae* cells were grown in carbon source-free synthetic medium (Sigma) supplemented with 5 g/L uniform ^13^C glucose (Cambridge isotope laboratories) and incubated overnight in the rotational shaker at 200 rpm and 30°C temperature. For the CIIST study of the *de novo* synthesis pathway activity, ^13^C microbial cells were pelleted down from the overnight ^13^C microbial culture using a centrifuge at 4000 rpm for 3 to 5 minutes. The pelleted cells were washed with PBS to remove the media trace. The ^13^C *E. coli* was reintroduced in the carbon source-free M9 minimal medium with ^12^C carbon source and incubated overnight in a rotational shaker at 180 rpm and 37°C. Similarly, ^13^C *S. cerevisiae* was also washed from ^13^C containing culture medium and reintroduced in the carbon source-free respective culture medium administered with ^12^C carbon source and incubated overnight in the rotational shaker at 200 rpm and 30°C incubation temperature. For the CIIST study of the *salvage pathway* activity, ^13^C microbial cells were pelleted down from the overnight ^13^C microbial culture via centrifugation at 4000 rpm for 3 to 5 minutes. The cell pellets were washed to remove the media trace using a centrifuge under the same parameters. For the control experiment, the ^13^C microbes were reintroduced in the carbon source-free M9 minimal medium with ^13^C glucose. For the salvage pathway monitoring, ^13^C microbes were reintroduced in the carbon source-free respective culture medium administered with ^13^C glucose supplemented with either purine (adenine and guanine) or pyrimidine (uracil and cytosine). The incubation of microbes was performed as per standard microbial culture protocols.

### Carbon isotope spectral tracing

Carbon isotope spectral tracing was done by obtaining Raman spectra from the multiple single *E. coli* and *S. cerevisiae* cells from different experimental conditions described above at increasing time intervals. First, cells were washed with PBS and streaked with proper dilution to get single cells on a clean Raman substrate. Raman spectra were acquired using an upright WITec confocal Raman spectrometer alpha 300 access equipped with 100x air objective, 532 nm laser and 600 gr/mm grating. Each acquired Raman spectrum was an average of 5 accumulations and 10-second laser exposure time. The raw Raman spectral data were preprocessed using MATLAB R2021B with Savitzky Golay smoothening and polynomial base fitting. Plots were generated using Origin Software.

### Carbon isotope imaging and analysis

For Carbon isotope imaging, Raman images of cells were collected from single *E. coli* and single *S. cerevisiae* cells grown in different experimental conditions described above at various time intervals. The cell pellets were washed and diluted to an optimised concentration to make sure single cells could be obtained before streaking on a clean Raman substrate. Raman image hyperspectral data was obtained using upright WITec confocal Raman spectrometer alpha 300 access equipped with 100x air objective, 532 nm laser and 600 gr/mm grating with 5 sec exposure time. The raw Raman hyperspectral data were pre-processed and the Raman images targeting ^12^C and ^13^C purine and pyrimidine representative peak position were generated using MATLAB 2021B software. Plots related to Raman were generated using Origin Software.

### Genomic DNA isolation and gel electrophoresis

^13^C *E. coli* and ^12^C *E. coli* cells were grown in the 20 ml culture medium. The cells were harvested by centrifugation at 7000 rpm for 10 mins at 4°C. The cells were resuspended in 1 ml Tris-EDTA buffer. The resuspended cells were taken into a 50 ml centrifuge tube and 1 ml SET (Sucrose EDTA and Tris) buffer with lysozyme and Proteinase K (400 μl) and incubated at room temperature for 90 minutes, lysate cleared up considerably. 20% SDS was added and shaken gently followed by 5 M NaCl and shaken gently. RNase was added, incubated for 5 minutes and shaken gently. Standard phenol-chloroform extraction was done 5 times with phenol-chloroform-isoamyl alcohol (25:24:1, pH 8.0). The aqueous layer was taken and chilled isopropyl alcohol was added and incubated at room temperature for 15 minutes followed by incubation at -20°Cfor 20 mins. Centrifugation was done at 13000 rpm for 20 minutes. The alcohol layer was decanted, pellets were washed with chilled ethanol and centrifugation was done at 13000 rpm for 5 minutes. Ethanol was discarded and the pellets were dried to remove the residual traces of alcohol. The pellet was reconstituted in double distilled DNase-free water in the microcentrifuge tube. Quantification was done using a nanodrop spectrophotometer. The integrity of the isolated genomic DNA (gDNA) was observed through agarose gel electrophoresis. Raman spectra were also acquired from the extracted gDNA.

## Results and Discussion

### Carbon isotope imaging reveals the spatial distribution of *de novo* synthesized purine and pyrimidine in single cells

Nitrogenous bases, classified as purine and pyrimidine play a crucial role in cellular metabolism. The key components of the *de novo* purine and pyrimidine synthesis pathways originate from carbon sources. The carbon flow from the source to purine and pyrimidine in these pathways has been represented in a schematic diagram **(Figures S1 A and S3 A)**. In order to non destructively monitor the newly synthesised purine and pyrimidine dynamic turnover in a single cell, *E. coli* was used as our prokaryotic model and *S. cerevisiae* was used as our eukaryotic model. CIIST methodology uses two isotopes of carbon as metabolite spectral tracers **(Figure 1A)**. The detection of guanine and adenine peak at around ∼1529 cm^-1^ (^13^C position) and ∼1576 cm^-1^ (^12^C position) while the uracil and cytosine at around ∼770 cm^-1^ (^13^C position) and ∼783 cm^-1^(^12^C position) were chosen as representatives spectral tracker for ^13^C and ^12^C purine and pyrimidine nitrogenous bases, **Table 1**.[13, 15, 17, 18, 21]

**Table 1.** Representative purine and pyrimidine representative ^12^C and ^13^C peak assignment as per previously reported studies.

| Sl. No. | <sup>12</sup>C position | <sup>13</sup>C position | Peak Assignment | Ref. |
| --- | --- | --- | --- | --- |
| 1. | 1576 | 1528 | Guanine, adenine (ring stretch) | [15] |
|  | 783 | 768 | Cytosine, uracil (ring breath) |  |
| 2. | 1578 | 1531 | Ring stretching of guanine and adenine | [18] |
|  | 787 | 770 | Cytosine, uracil |  |
| 3. | 1574 | 1527 | Nucleic acid | [21] |
|  | 783 | 769 | Cytosine, uracil (C=O, C-N and ring deformation) |  |
| 4. | 1574 | 1529 | Guanine, adenine (ring stretching) | [13] |
|  | 785 | 769 | Cytosine, uracil |  |
| 5. | 785 | 770 | Cytosine, uracil (ring stretch) | [17] |

**Figure 1.**
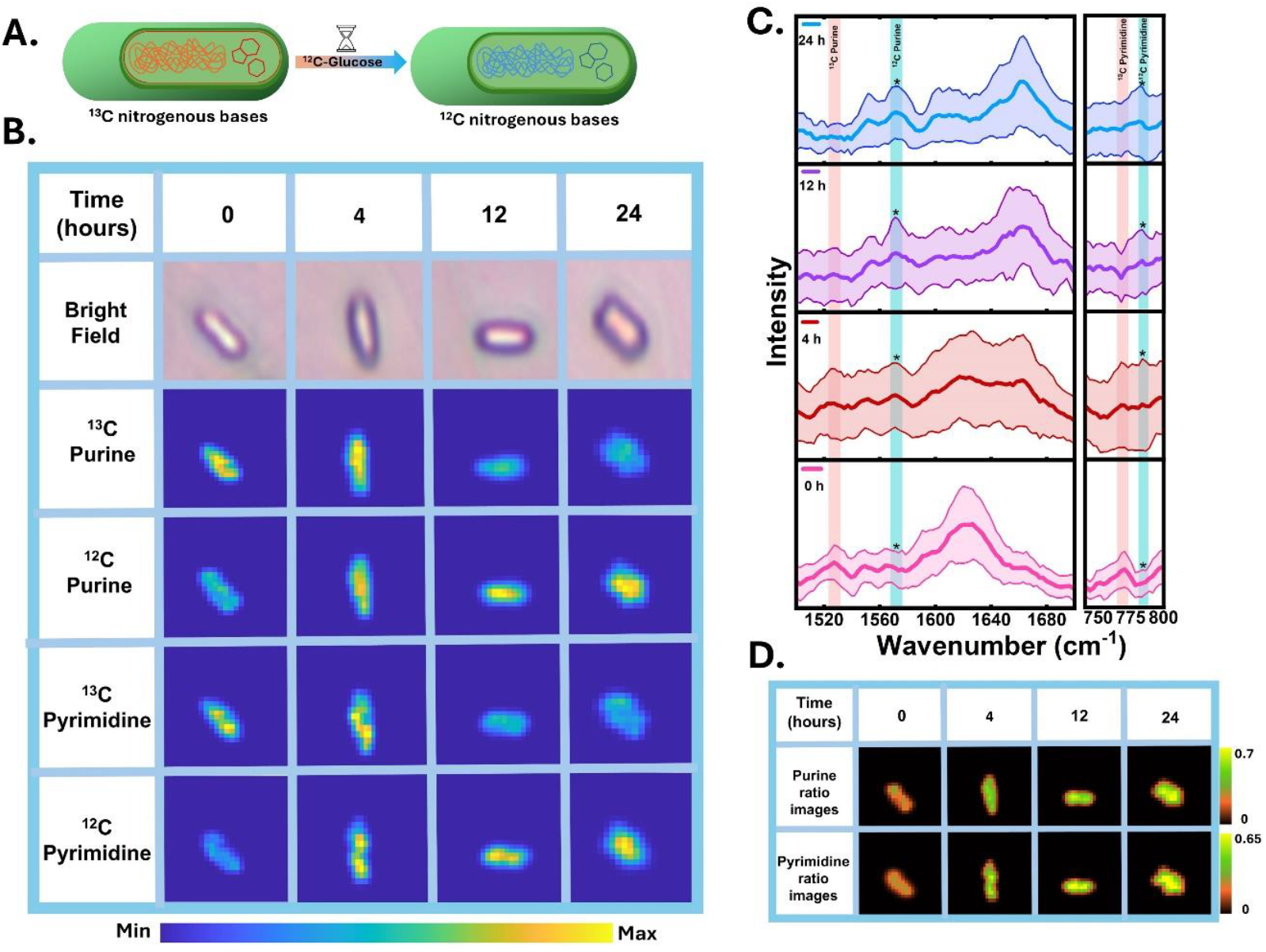
Visualising of the *de novo* synthesized nitrogenous bases in *E. coli* by carbon isotope imaging. (A)Schematic showing carbon isotope imaging and spectral tracing methodology used for monitoring synthesis of nitrogenous bases in ^13^C cell grown in ^12^C carbon source producing ^12^C nitrogenous bases. (B)Raman images of a single *E. coli* cell at different incubation time points targeting the ^13^C and ^12^C position of purine and pyrimidine (C) Average Raman spectra from pixels under the Raman image boundary of *E. coli* cells showing the blue shift of ^13^C and ^12^C nitrogenous base representative peak. (standard deviation is shown as the shaded area) (C) Purine and pyrimidine ratio images between the Raman image at ^12^C nitrogenous base and total nitrogenous base (^12^C+^13^C), representing the newly synthesized nitrogenous base turnover at multiple time intervals.

Hyperspectral imaging was performed to visualize the spatial distribution of the *de novo* synthesised nitrogenous bases and their turnover dynamics in a single cell **(Figure 1B)**. The carbon isotope images were generated at 0, 4, 12 and 24 hours post incubation. At 0 hr, the carbon isotope image of the *E. coli* cells shows high-intensity spatial distribution at ^13^C isotope position, primarily localised in the cytoplasmic region where *de novo* nitrogenous base synthesis occurs and nitrogenous bases derived metabolites as well as nucleoid predominates. In contrast, the ^12^C position shows lower intensity indicating the initial existing pool of the nitrogenous bases in the inoculate cell is ^13^C isotope enriched. After 4 hours of incubation time, interestingly we observed a similar intensity spatial distribution from the ^13^C as well as ^12^C position. This indicates the co-existence of already existing and *de novo* newly synthesized nitrogenous base pools in the cell. However, at latter time points 12 and 24 hours the distribution pattern completely shifts when compared to cells at 0 hours: the ^13^C intensity spatial distribution is seen to be diminished drastically, while the ^12^C intensity distribution becomes highly enhanced. This enhanced ^12^C position distribution pattern clearly shows the nitrogenous base *de novo* synthesis pathway in action resulting in newly synthesized nitrogenous bases gradually replacing the existing pool over time. In addition to being capable of imaging the *de novo* synthesized new nitrogenous bases distribution in a single cell, we further validated our findings by spectral tracing in the mean Raman spectra along with its standard deviation from all pixels within the cellular boundary of the Raman image at different time intervals, which revealed the blue shift pattern. **(Figure 1C)** The standard deviation from ^13^C as well as the ^12^C position also gives meaningful information about the spatial distribution pattern that the nitrogenous bases are distributed heterogeneously within the cells.

Further, we presented a more quantitative perspective of carbon isotope imaging for visualizing the newly synthesized nitrogenous base turnover dynamics by generating a ratio carbon isotope image between the newly synthesized nitrogenous bases and the total nitrogenous base (either purine or pyrimidine, **Figure 1D**. The ratiometric intensity of the hyperspectral data were calculated and mapped by formulae:

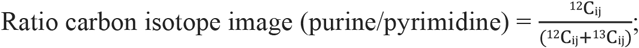

where i and j are the pixel positions, ^12^C_ij_ is the intensity value of the ^12^C peak position at each pixel and ^13^C_ij_ is the intensity value of the ^13^C peak position at each pixel distributed over the carbon isotope image of different time points. The ratio carbon isotope image depicts the progressive ratiometric intensity change with time (**Figure 1D**). The increasing ratiometric intensity distribution at different time intervals semi-quantitatively demonstrates the ^12^C assimilation and turnover dynamics of newly synthesized nitrogenous bases in cells.

Previous studies have well documented the peak positions of ^13^C and ^12^C isotope purine and pyrimidine in cells with vibrational mode, as presented in Table 1. We further validated the incorporation of the newly synthesized nitrogenous bases in the cellular macromolecules and the corresponding peaks. We extracted the DNA from the ^13^C and ^12^C *E. coli* cells and performed the agarose gel electrophoresis test (**Figure S2A**). The genomic DNA was observed to be positioned near the top of the gel above the DNA ladder marker indicating an unfragmented large size. Before obtaining Raman spectra, DNA yield and purity were also measured by UV absorption. The obtained ^13^C DNA yield was 603.2 ng/µl and A(260/280) was 1.91. The obtained ^12^C DNA yield was 882.2 ng/µl and A(260/280) was 1.86, so it met the experimental requirements. After this, the Raman spectra were recorded from extracted ^13^C DNA and ^12^C DNA and we got the representative peaks for ^12^C/^13^C purine and pyrimidine as we got in the whole single cells (**Figure S2B**). This validates that the characteristic Raman peaks observed for ^12^C/^13^C purine and pyrimidine in whole cells are indeed attributable to nitrogenous bases, and also confirms the successful spectral tracing of the incorporation of newly synthesized nitrogenous bases into the nucleic acid.

Similar experiments were performed in *S. cerevisiae*. The spatial distribution of these newly synthesized nitrogenous bases in yeast cells was mapped and the high-intensity distribution seems to be localised in the cytoplasmic region (**Figure 2A**). Similar, to the isotope images of *E*. *coli* cells, the yeast cells also demonstrate gradual shifts of the intensity spatial distribution over time from the ^13^C peak position towards the ^12^C peak position demonstrating the newly synthesized nitrogenous base in the single cell level. In addition to imaging the distribution of newly synthesized nitrogenous bases at the single-cell level, we validated our findings by spectral tracing the blue shift from the mean Raman spectra with standard deviation from pixels within the cellular boundary of cell image at different time points (**Figure 2B**). The nascent nitrogenous bases turnover dynamics in the yeast cell using hyperspectral imaging, we generated ratiometric Raman images of the yeast cell (newly synthesized nitrogenous bases divided by entire nitrogenous base pool, either purine or pyrimidine) using the same formulae mentioned earlier. These ratio images show a clear change in turnover dynamics over time (**Figure 2C**). The findings suggest that the CIIST method can have wide applicability in studying pathways of the broad band of culturable microbes and in non-destructive microbial metabolomics qualitatively and semi-quantitatively.

**Figure 2.**
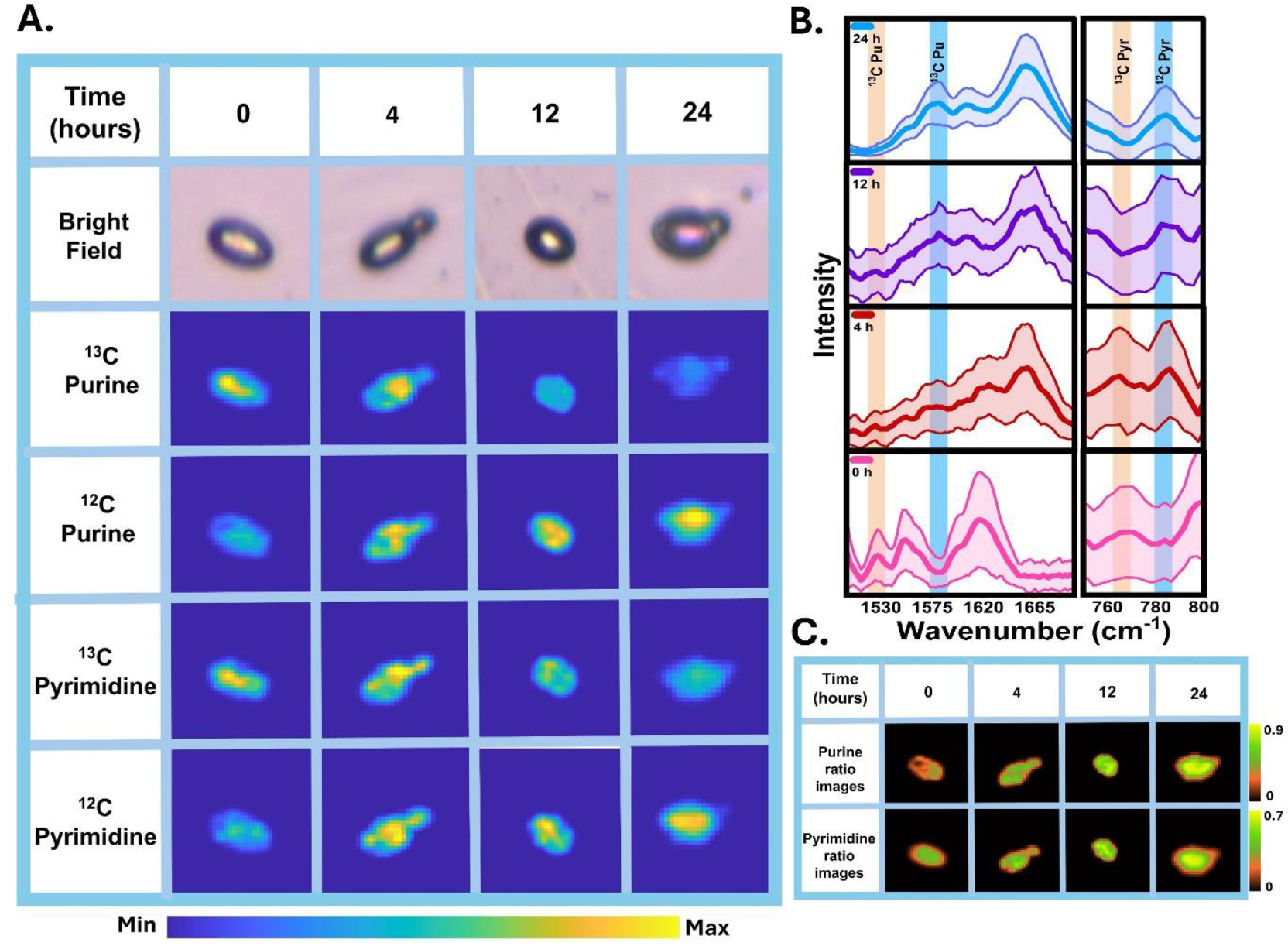
Visualising of the *de novo* synthesized nitrogenous bases in *S. cerevisiae* by carbon isotope imaging. (A) Raman images of a single *S. cerevisiae* cell at different incubation time points targeting the ^13^C and ^12^C position of purine and pyrimidine (C) Average Raman spectra from pixels under the Raman image boundary of *S. cerevisiae* cells showing the blue shift of ^13^C and ^12^C nitrogenous base representative peak. (standard deviation is shown as the shaded area) (C) Purine and pyrimidine ratio images between the Raman image at ^12^C nitrogenous base and total nitrogenous base (^12^C+^13^C), representing the newly synthesized nitrogenous bases turnover at multiple time intervals.

### Spectral tracing of the cell-to-cell metabolic heterogeneity of purine and pyrimidine *de novo* synthesis activity

In order to track the heterogeneity of the purine and pyrimidine *de novo* synthesis activity carbon isotope spectral tracing in multiple cells was performed. Raman spectra were acquired from multiple *E. coli* single cells and multiple *S. cerevisiae* single cells at different time intervals at 0, 2, 4, 8, 12 and 24 hours using point spectroscopy. The mean Raman spectra of *E. coli* cells (**Figure S1 B & C**) and multiple single *S. cerevisiae* cells (**Figure S3 B & C**) were analysed along with the standard deviation. At 0 hour, intensified Raman peak from representative purine and pyrimidine ^13^C position from cells was observed. This indicates that the total existing pool of nitrogenous bases is enriched with ^13^C isotope. After the cells were reintroduced in a culture medium with a ^12^C isotope as the sole carbon source, new peaks were detected emerging at ^12^C position along with the Raman peak at ^13^C position at 2 and 4 hours. This newly emerged peak at the ^12^C position indicates the *de novo* synthesis nitrogenous bases activity. At later time intervals - 8, 12 and 24 hours, an intensified signal at the ^12^C position was observed. This time-dependent blue shift of peak from ^13^C position towards ^12^C position in representative peaks of nitrogenous bases gives insight into the gradual carbon isotope integration and turnover of *de novo* synthesized new nitrogenous base into the already existing pool in multiple microbial single cells, qualitatively and standard deviation shows cells metabolic heterogeneity. For the semi-quantitative analysis of ^13^C incorporation, we studied the varying intensity ratiometric (^12^C/^13^C purine and pyrimidine) at different time intervals as shown in the line plot for *E. coli* cells (**Figure S1 D & E**) and S. cerevisiae cells (**Figure S3 D & E**). The increasing ratiometric intensity at different time intervals semi-quantitatively demonstrates the ^12^C isotope assimilation and turnover dynamics of newly synthesized nitrogenous bases in multiple single cells.

### Carbon isotope imaging visualizes salvage pathway activity in single cells and *in situ* peak validation

The cells can acquire purine and pyrimidines exogenously from the extra-cellular environment or recycle nitrogenous bases and their derivatives through the salvage pathway. The schematic diagram of the salvage pathway in *E. coli* (**Figures S1 A**) and *S. cerevisiae* (**Figures S3 A**) shows the crosslinks between different purines and pyrimidines. For visualizing the salvage pathway activity using CIIST, the first challenge was to separate the internal nitrogenous bases signal from the exogenous nitrogenous bases from the extracellular environment three experimental conditions were performed (i) ^13^C cells were grown in the ^13^C glucose as the only carbon source in the culture medium (Figure 3A(i)), (ii) ^13^C cells were grown in the ^13^C glucose as the only carbon source in the culture medium with ^12^C purine as exogenous purine base (Figure 3A(ii)), (iii) ^13^C cells were grown in the ^13^C glucose as the only carbon source in the culture medium with ^12^C purine as exogenous pyrimidine base(Figure 3A(iii)). The hyperspectral images were acquired from single cells at 0 and 12 hours with three experimental conditions for *E. coli* (Figure 3B) and *S. cerevisiae* (Figure 4A). At zero hour, we observe that intensity spatial distribution is higher for the ^13^C position and lower for the ^12^C position. This clearly indicates the total pool at 0 hour is ^13^C enriched. At 12 hours (i), we observe that the signal intensity spatial distribution is similar to that of 0 hours with higher intensity at the ^13^C position as there is no exogenous purine or pyrimidine source in the extracellular environment. At 12 hours (ii), we observe that the signal intensity spatial distribution is similar to that of 0 hours with intensity distribution at the ^13^C position of both purine and pyrimidine. However, the ^12^C position of purine is also seen to have a higher signal intensity distribution in the cytoplasmic region of both *E. coli* and *S. cerevisiae* whereas the signal intensity distribution from the ^12^C position of pyrimidine has a low-intensity distribution. This result shows the purine salvage activity by monitoring the incorporation of exogenous purine from the extracellular environment into the cell. Similarly at 12 hours (iii), along with intensity distribution from ^13^C position, a higher signal intensity distribution for ^12^C pyrimidine was also observed in the cytoplasmic region of both *E. coli* and *S. cerevisiae*. This hyperspectral image shows the pyrimidine salvage activity by monitoring the incorporation of exogenous pyrimidine from the extracellular environment into the cell at the single-cell level. Taking our analysis a step forward, we further confirmed our findings by carbon isotope spectral tracing by analyzing the mean Raman spectra with standard deviation from the pixels within the cellular boundary for *E. coli* (**Figure 3C**) and *S. cerevisiae* (**Figure 4B**). Additionally, we also examined the ratiometric intensity variations (^12^C/^13^C purine and pyrimidine Raman peaks) over different time intervals, as shown in the line plots (Figure 4E). The ratiometric intensity only changes for the cells supplemented with either ^12^C purine (Figure S4 A/B – 12(ii)) or ^12^C pyrimidine((Figure S4 C/D – 12(iii)) which provides a semi-quantitative representation of metabolic exogenous assimilation of purine and pyrimidine in the cell.

**Figure 3.**
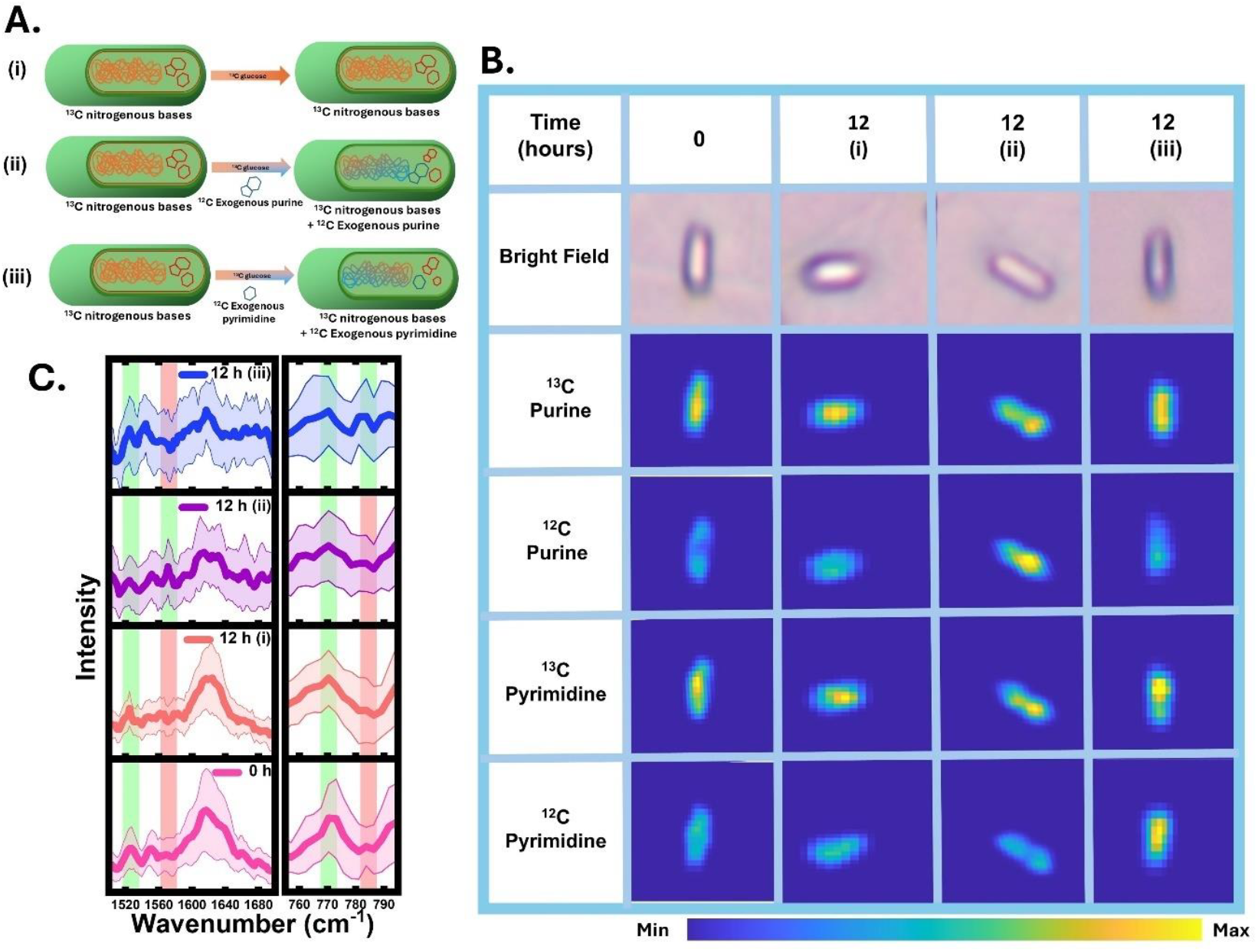
Carbon isotope imaging for visualising exogenous nitrogenous bases in *E. coli*. (A) Schematic diagram showing all three experimental conditions for CIIST: (i) ^13^C cells grown in ^13^C glucose as only carbon source (ii) ^13^C cells grown in ^13^C glucose supplemented with ^12^C purine in culture medium (iii) ^13^C cells grown in ^13^C glucose supplemented with ^12^C pyrimidine culture medium. (B) Visualising of the new nitrogenous bases incorporated into the *E. coli* cell exogenously using carbon isotope imaging at 0 and 12 hours: (i) control (ii) purine salvage pathway (iii) pyrimidine salvage pathway. (C) Average Raman spectra from all the pixels under the Raman image boundary of *E. coli* cell at 0 and 12 hours at all three experimental conditions (standard deviation is shown in shaded region

**Figure 4.**
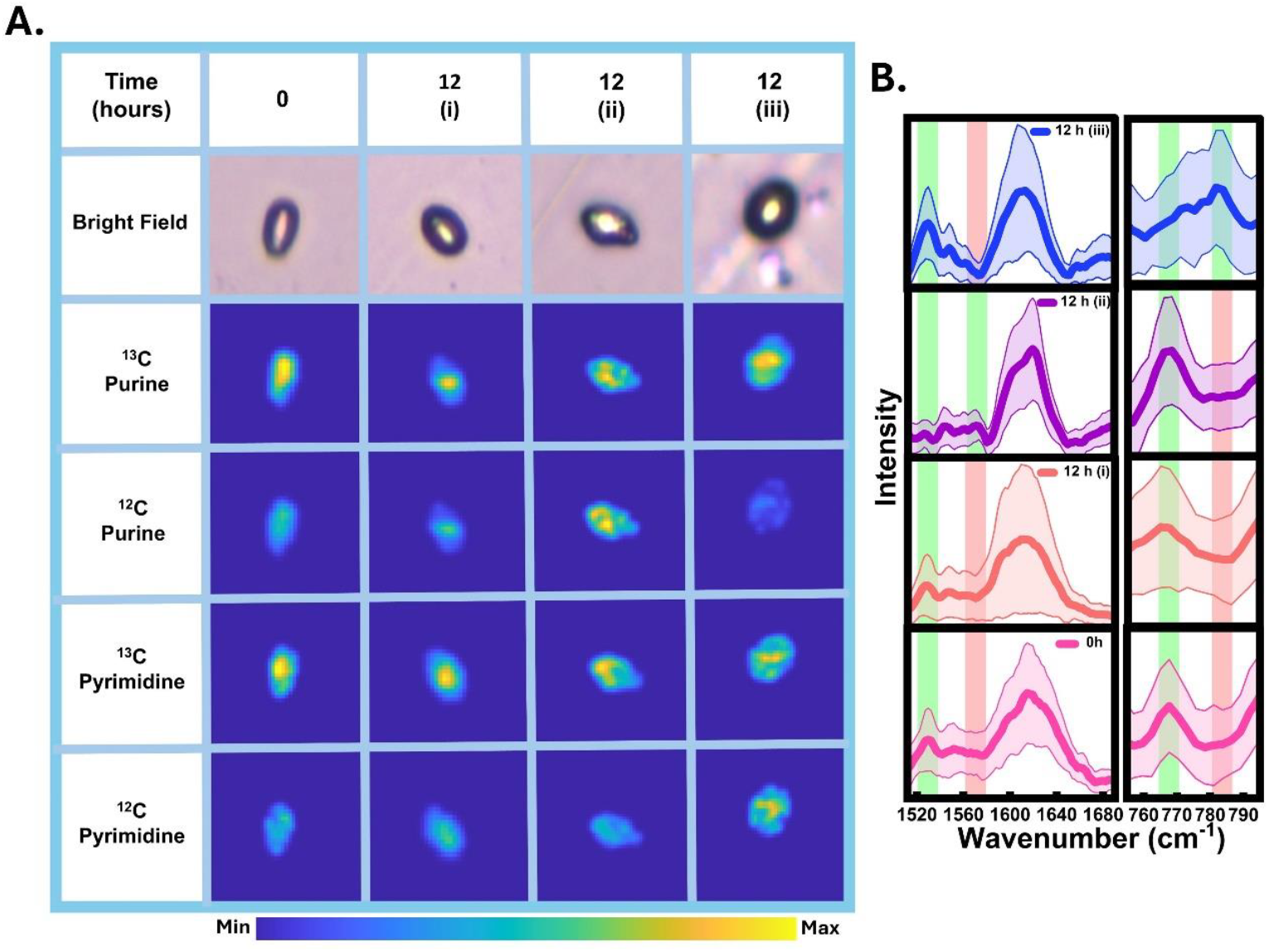
Carbon isotope imaging for visualising exogenous nitrogenous bases in *S. cerevisiae*. Visualising of the new nitrogenous bases incorporated into the *S. cerevisiae* cell exogenously using carbon isotope imaging at 0 and 12 hours: (i) control (ii) purine salvage pathway (iii) pyrimidine salvage pathway. (C) Average Raman spectra from all the pixels under the Raman image boundary of *S. cerevisiae* cell at 0 and 12 hours at all three experimental conditions (standard deviation is shown in shaded region.

In this study, we visualize the purine and pyrimidine *de novo* and salvage pathway metabolic activity of purines and pyrimidines using CIIST methodology. Our findings opens a new adjuct methodology for metabolic mapping at single cell level which can provide spatial and temporal maps offering new insight into nucleotide metabolism. Carbon isotope imaging explores the potential of spontaneous confocal Raman imaging based on point-per-line and lines-per-image. Our results indicate a high concentration of nitrogenous bases localized within the cytoplasmic regions of the cells, detected without the use of molecular tags or labels. Since the pyrimidine representative peak includes contributions from both uracil and cytosine, future studies with higher-resolution CIIST could potentially enable the precise localization and differentiation of nucleic acids, such as RNA and DNA, at the single-cell level. This advancement would further enhance the capability of hyperspectral imaging in metabolic mapping for advancing genomics and transcriptomics. Previously, Chisanga et al combined carbon isotope with point spectroscopy to study the carbon isotope incorporation kinetics in the *P. putida* targeting nitrogenous base peak (766 → 783 cm^−1^). They reported the faster assimilation of carbon isotope into nitrogenous bases and amides than phenylalanine.[15] Muhamadali et al. reported using point spectroscopy that in *E. coli* cells enriched with ^13^C, the corresponding peak assigned to the stretching vibration of cytosine and uracil appears at 769 cm^-1^ whereas when it is ^15^N enriched it appears at 778 cm^-1^. Additionally, they also reported 47 cm^-1^ wavenumber position red shift for ^13^C enriched cells and 8 cm^-1^ wavenumber redshift for ^15^N enriched cell compared to the ^12^C/^14^N enriched peak position at 1574 cm^-1^.[21] Although our study explores nitrogenous base metabolism, we use carbon isotope enrichment instead of nitrogen isotope enrichment in cells because carbon isotopes produce a larger spectral shift. This results in better peak resolution, making it more suitable for hyperspectral imaging and enhancing the robustness of CIIST. In previous studies, carbon isotope has been widely used to track the carbon flow using point spectroscopy as it provides precise peak identification and faster detection of peak shift due to isotopic enrichment. Moreover, the intensity ratio of the heavier to lighter isotope or vice versa can be used as a semi-quantitative scale to measure metabolic changes. However, despite these advantages, point spectroscopy is limited in its spatial coverage, as it collects data from a single point. This makes large-area scanning methods such as CIIST a crucial technique for dynamic metabolic study. CIIST based on Raman imaging contains both spatial and spectral information enhancing its applicability for high-throughput analysis. Barzan et al. have also generated a chemical map of bacteria targeting 780 cm^-1^ in air-dried and freeze-dried cells.[22] Their chemical map exhibits a similar distribution to the one generated in our results; however, our approach provides additional insights by capturing information on newly synthesized metabolites, such as purines and pyrimidines. This highlights the advantage of our method in tracking dynamic metabolic processes with greater specificity. As Raman scattering is a weak phenomenon the spatial map we generated has a limiting factor due to the trade-off between spectral acquisition time and spatial coverage.[23–25] One of the significant limitations of this study is the time required for acquiring hyperspectral imaging data from cells. Increasing the step size and minimizing the laser exposure time can be one way to address it; however, it may result in undersampling and a lower signal-to-noise ratio, ultimately decreasing spectral quality and spatial resolution. Improving the spatial resolution is another challenge with spontaneous Raman imaging in microbial cells which needs to be addressed in future. Stimulated Raman spectroscopy imaging combined with CIIST can be a future milestone in the field of metabolic mapping which has been explored previously using deuterium isotopes in both microbes and mammalian cells.[26–28] Although limitations with resolutions are there; however, CIIST is a non-destructive and minimally invasive approach which has the potential applicability to study nucleotide metabolism in animal cells also *in situ*. Our study shed an experimental light on multiplex imaging of purine and pyrimidine metabolic pathway activity taking hyperspectral imaging as another promising complementary tool for nitrogenous base metabolism.

## Conclusion

This study demonstrates the successful application of Carbon Isotope Imaging and Spectral Tracing(CIIST) for visualizing the *de novo* and salvage pathway activity of nitrogenous bases in single microbial cells (prokaryotes as well as eukaryotes). By tracing the shift due to the carbon isotopic effect using point Raman spectroscopy, we successfully performed a comprehensive spectral tracing study, qualitatively and quasi-quantitatively. This study provides insights into the metabolic turnover of newly synthesized nitrogenous base via *de novo* pathway and exogenous nitrogenous base assimilation via the salvage pathway *in situ*. This work highlights carbon isotope imaging and spectral tracing “CIIST” as a powerful adjunct metabolomics tool for non-destructive, real-time nitrogenous base pathway mapping in single-cell systems, advancing our understanding of the complex spatial and temporal dynamics in situ.

## Availability of data and materials

The datasets used and/or analysed during the current study are available from the corresponding author on reasonable request.

## Acknowledgements

This work was carried out under research grant project no (37/1739/23/EMR-II) supported by the Council of Scientific and Industrial Research (CSIR), Government of India and project no. IIRP-2023-1734 from the Indian Council of Medical Research (ICMR), Government of India.

## Supplementary Information

**Figure S1.**
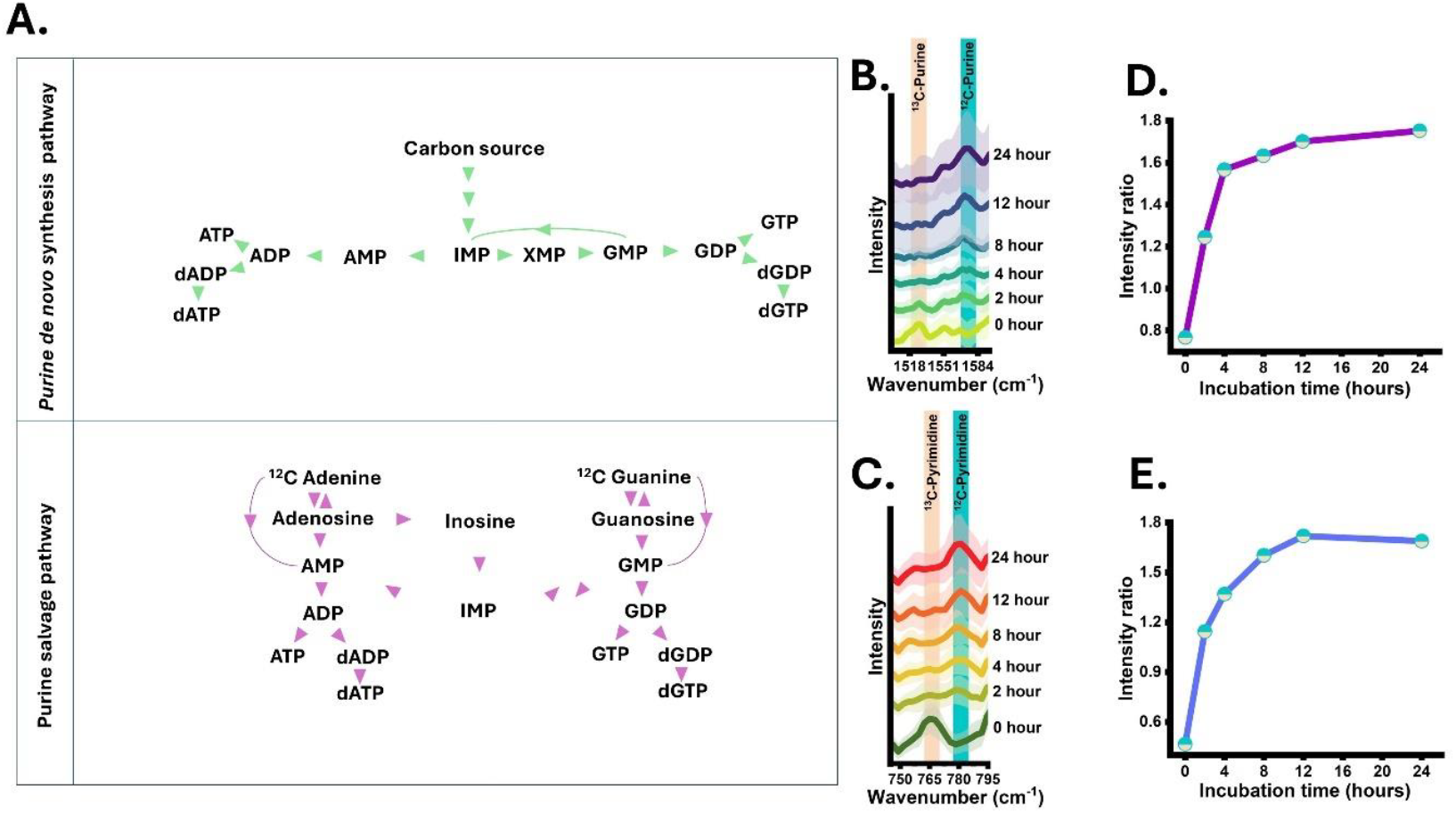
(A) Schematic representation of the purine *de novo* and salvage pathways in *E. coli*. (B) Average Raman spectra of multiple *E. coli* cells at different incubation time points showing *de novo* synthesis of purines (standard deviation is shown as shaded area) (C) Average Raman spectra of multiple *E. coli* cells at different incubation time points showing *de novo* synthesis of pyrimidine (standard deviation is shown as shaded area) (D) Intensity ratio plot (^12^C purine/ ^13^C purine) showing turnover of newly synthesized purine to the old pool of purine at different incubation time points. (E) Intensity ratio plot (^12^C pyrimidine/ ^13^C pyrimidine) showing turnover of newly synthesized pyrimidine to old pool of pyrimidine at different incubation time points. (standard deviation is shown as shaded area)

**Figure S2.**
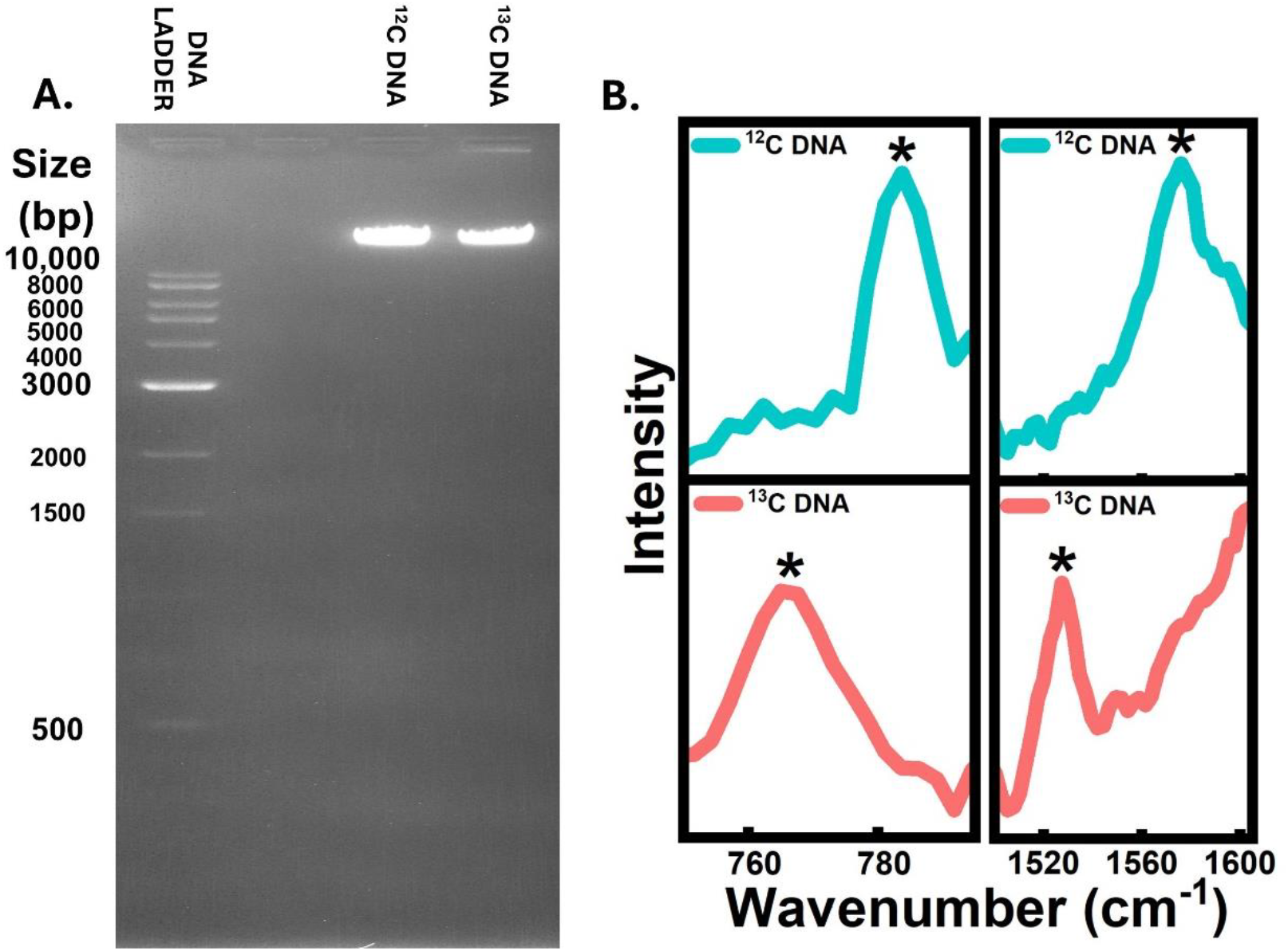
gDNA extracted from ^13^C *E. coli* and ^12^C *E. coli*. (A) Agarose gel electrophoresis of ^13^C and ^12^C DNA with DNA ladder lane. (B) Raman spectra of ^13^C and ^12^C extracted DNA show shifted representative peak positions for purine and pyrimidine nitrogenous bases.

**Figure S3.**
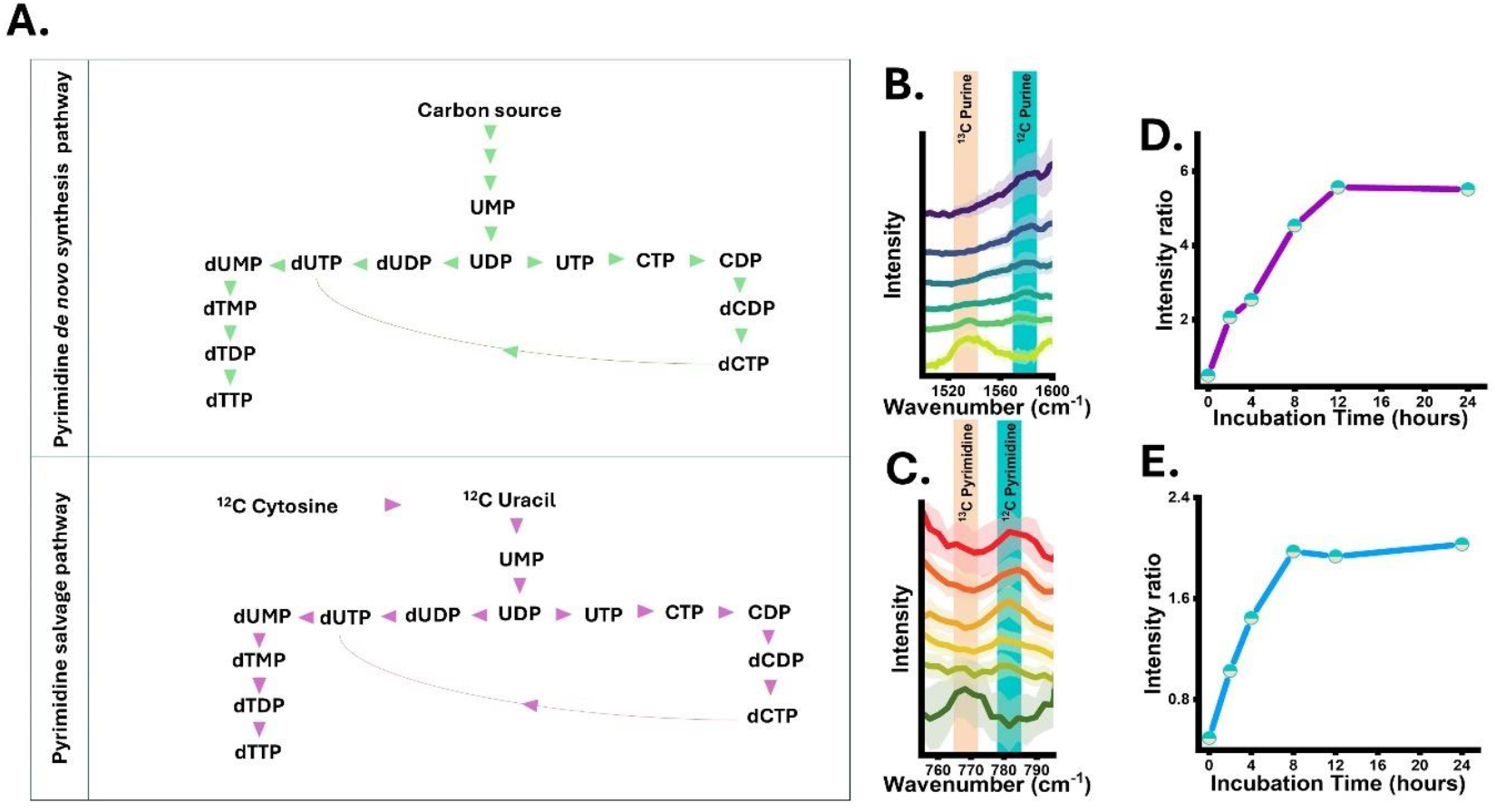
(A) Schematic representation of the purine *de novo* and salvage pathways in *S. cerevisiae*. (B) Average Raman spectra of multiple *S. cerevisiae* cells at different incubation time points showing *de novo* synthesis of purines (standard deviation is shown as shaded area) (C) Average Raman spectra of multiple *S. cerevisiae* cells at different incubation time points showing *de novo* synthesis of pyrimidine (standard deviation is shown as shaded area) (D) Intensity ratio plot (^12^C purine/ ^13^C purine) showing turnover of newly synthesized purine to the old pool of purine at different incubation time points. (E) Intensity ratio plot (^12^C pyrimidine/ ^13^C pyrimidine) showing turnover of newly synthesized pyrimidine to old pool of pyrimidine at different incubation time points. (standard deviation is shown as shaded area)

**Figure S4.**
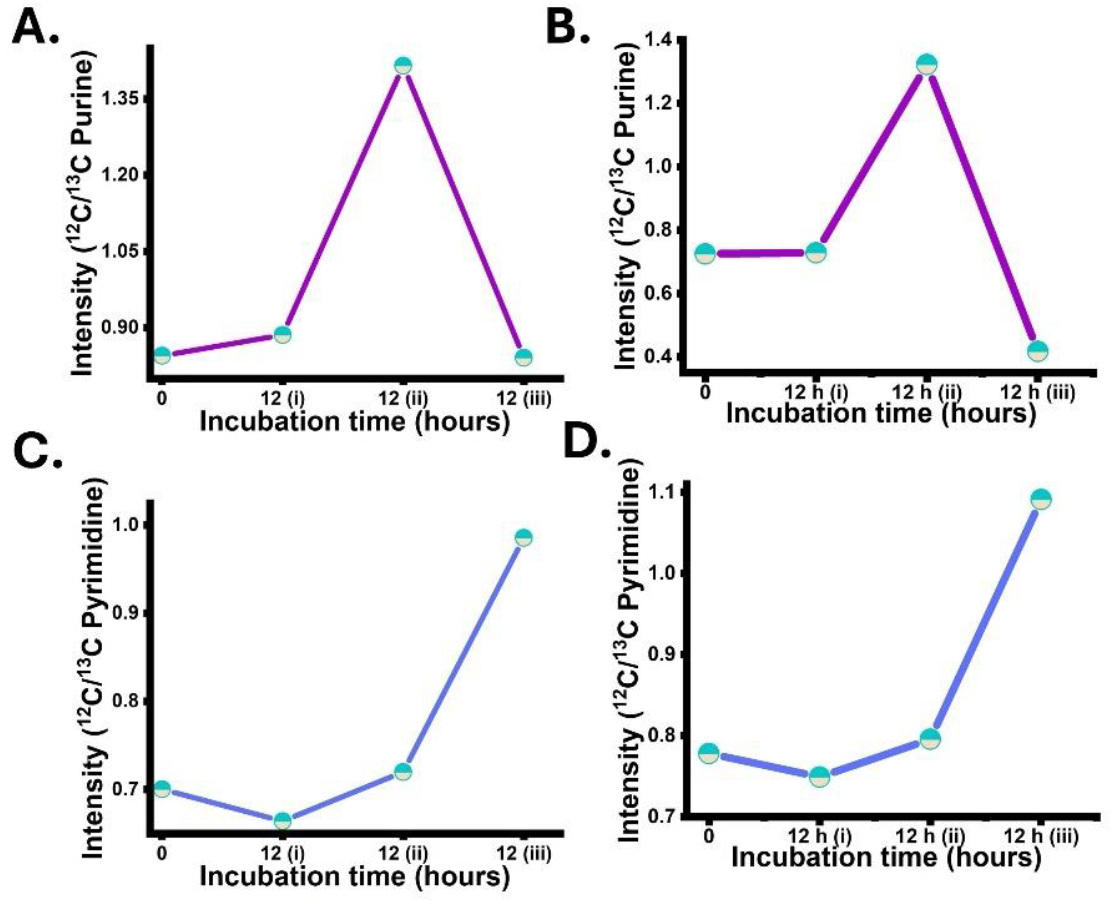
Ratiometric intensity plot for representative peak from hyperspectral image *E*.*coli* (A & C) and *S. cerevisiae* (B & D) at 0 and 12 hours with three experimental conditions.

